# Peripheral Complement C4 Protein in Schizophrenia: Association with Gene Copy Number and Immune Cell Subtypes

**DOI:** 10.1101/2025.09.16.676439

**Authors:** Agnieszka Kalinowski, Claudia Macaubas, Lauren Anker, Diane E. Wakeham, Marcus Ho, Reenal Pattni, Batuhan Bayram, Surbhi Sharma, Joanna Liliental, Jong H. Yoon, Elizabeth D. Mellins, Alexander E. Urban

**Affiliations:** Department of Psychiatry and Behavioral Sciences, Stanford University School of Medicine, Stanford, CA; Department of Pediatrics, Stanford University School of Medicine, Stanford, CA; Kaiser Permanente, San Francisco, CA; Department of Genetics, Stanford University School of Medicine, Stanford, CA; Department of Medicine, Division of Research and Education in Academic Medicine (DREAM), Stanford University School of Medicine, Stanford, CA; Translational Applications Service Center (TASC), Stanford University School of Medicine, Stanford, CA; Translational Research and Applied Medicine Center (TRAM), Stanford University School of Medicine, Stanford, CA; Palo Alto Veterans Healthcare System, Palo Alto, CA

**Author notes:** co-corresponding authors: Alexander E. Urban,3165 Porter Drive, Palo Alto, CA 94305, Agnieszka Kalinowski, 3165 Porter Drive, Palo Alto, CA, 94305. deceased March 24, 2024 (Elizabeth D. Mellins).

## Abstract

The lack of disease-modifying treatments for schizophrenia necessitates the exploration of novel aspects of its pathophysiology, including innate immune mechanisms in the periphery. C4 protein activation, associated with the complement cascade of innate immunity, associates with symptoms and predicts outcomes. However, C4 protein activation does not coincide with expected changes to other proteins in the complement cascade, suggesting another source of C4 protein activation. Using a combination of fresh whole blood from ten anonymous donors and a large set of publicly available microarray data, we show, for the first time, that C4 protein is found and expressed primarily in neutrophils and monocytes. Then, we compared the correlation between C4 protein in neutrophils, classical monocytes, plasma and the number of C4A gene copies. We determined the number of C4A genes using digital droplet PCR, C4 protein in neutrophils (15 patients/21 controls) and plasma (30 patients/38 controls) using western blotting, and classical monocytes (30 patients/38 controls) using flow cytometry. We found a moderate positive correlation between the number of C4A gene copies and the amount of C4 protein *only* in neutrophils and *only* in the schizophrenia group (Spearman’s rho, r = 0.64, p = 0.01, Cohen’s d ∼1.67). Our results indicate a convergence of innate immunity mechanisms associated with schizophrenia. This novel mechanism of innate immunity in schizophrenia deserves further study to determine whether it could be a useful drug target.

## Introduction

Given the lack of disease-modifying agents to treat schizophrenia (SCZ), understanding its pathophysiology is imperative. Convergent evidence from epidemiological, animal, and clinical studies points to innate immune system activation, often referred to as the *innate immune hypothesis,* as an important element of the disease mechanism (1–3). Large epidemiological studies have shown that severe infections in early childhood that stimulate the innate immune response are significant risk factors for SCZ (4,5). The number of neutrophils and monocytes, which are cells of the innate immune system, are increased in people experiencing their first episode of psychosis (FEP) and chronic SCZ (6). Innate immune activators, such as cytokines and proteins involved in the complement cascade, are elevated in both the serum and cerebrospinal fluid (CSF) of individuals with chronic SCZ, FEP, and in the serum of individuals at clinical high risk (CHR) for psychosis (3,7–12). These circulatory immune factors have been associated with neuroimaging findings, symptoms, and cognitive measures in SCZ (13,14), and predict clinical outcomes in CHR and SCZ (15,16).

Understanding the key pathological mechanisms of innate immunity in SCZ may facilitate identification of novel therapeutic targets. The relevance of peripheral immune mechanisms to brain functioning has been demonstrated (17,18). And modifying the peripheral circulation improves cognition in Alzheimer’s disease in animal and human studies (19,20). However, clinical trials investigating drugs targeting the innate immune response in schizophrenia have yielded inconsistent results (21–23). These mixed outcomes underscore the need to better understand how the innate immune system is altered in SCZ.

A consistent finding of complement activation in individuals at risk for and with SCZ, specifically involving C4 protein, offers a promising line of investigation (10,11,15,24). C4 protein is expressed by the C4A and C4B genes. The number of C4A gene copies is associated with the risk of SCZ in genome-wide association studies (GWAS) of European ancestry (25). Higher C4A gene expression has been associated with higher levels of synaptic pruning in animal and *in vitro* induced-pluripotent stem cell studies (26,27), suggesting a link between C4A gene copies and excessive synaptic pruning. This *excessive pruning* hypothesis is further supported by the finding of higher C4A gene expression in postmortem brain tissue and elevated C4A protein in cerebrospinal fluid in individuals in FEP who develop SCZ (25,28,29). While the evidence points to C4A gene mechanisms in the brain, emerging data suggests that they may also be important in the periphery (10,11,24,30). In plasma, C4 protein is part of the classical or lectin pathway of the complement cascade, an arm of innate immunity (24). Upon activation, C4 protein is enzymatically cleaved to produce an activation product, C4-anaphylotoxin (C4-ana) and a cleaved C4 protein that binds other proteins in the complement cascade. Measuring the concentrations of activation products and cleaved proteins along parts of the complement cascade allows us to map which portion of the complement cascade is active. SCZ clinical studies consistently find increased concentrations of C4-ana and decreased concentrations of its inhibitor, C4-BP, (which suggests high utilization) in plasma samples from individuals at risk for and with SCZ (11,12,15,24,30). However, clinical studies have not found consistent evidence of a corresponding decrease in C4 protein concentration or changes in the concentrations of accompanying proteins along a specific pathway of the complement cascade in SCZ samples (30–32). This inconsistency suggests that C4 protein is responsible for the increased concentration of C4-ana in SCZ samples, which may originate from a non-plasma source in the peripheral circulation. Complement proteins in the plasma are produced in the liver (33,34). However, recent findings have revealed that some complement proteins are both expressed and activated inside distinct immune cell types (35). This led us to ask whether (1) C4 protein could be produced and activated in/on distinct immune cell types (neutrophils and monocytes), and (2) whether the amount of C4 protein in neutrophils and/or monocytes is decreased because of C4 protein activation in SCZ samples compared to controls.

To test our hypothesis that the C4 protein in neutrophils and monocytes is activated in SCZ, we conducted a two-part study. In part one, to establish whether C4 protein is expressed in neutrophils and/or monocytes, we used publicly available gene expression data and validated which immune cells contain C4 protein using whole blood samples from a small cohort of anonymous blood donors. These studies allowed us to confirm, for the first time, that C4 protein is expressed and present in neutrophils and monocytes. In the second part of our study, we compared the amount of C4 protein (as a function of the number of C4A gene copies) in neutrophil and monocyte samples from individuals with early SCZ and controls. We expected C4 protein to be expressed and consumed via C4 protein activation in the setting of innate immune system activation in SCZ. Therefore, we reasoned that examining the correlation between cell-associated C4 protein and the number of C4A gene copies may more accurately reflect the dynamic process of active C4 protein expression and activation/consumption. Similarly, in our previous study, we found a preliminary association between C4 protein activation (plasma C4-ana concentration) and the number of C4A gene copies (24). Finally, we conducted exploratory analyses comparing the amount of C4 protein in different immune cell populations in samples from SCZ and controls.

## Methods

### I: Unbiased examination of C4 protein in immune cell types using fresh whole blood

To determine which immune cell types contain C4 protein in an unbiased manner, we collected fresh whole blood samples from volunteers who donated blood to the Stanford Blood Center on the day we performed the experiment. Whole blood was collected by venipuncture in 6 mL K2 EDTA blood collection tubes and kept at room temperature. Whole blood samples were stained with antibodies against the major immune cell subtypes: CD3, CD14, CD16, CD19, CD45, CD56, CD66b and HLA-DR (Supplementary Table 2 contains details of flow cytometry antibodies used). Red blood cells were then lysed, and white blood cells were fixed using the eBiosciences 1-step Fix/Lyse solution (Invitrogen, Waltham, MA) according to the manufacturer’s instructions. White blood cells were collected by centrifugation (500 × g for 5 min at room temperature). To access both surface and intracellular C4 protein, we permeabilized white blood cells by resuspending the centrifuged cell pellet in Permeabilization Buffer according to manufacturer instructions (eBioscience Permeabilization Buffer, Invitrogen). Then, cells were resuspended in permeabilization buffer and an antibody against the C4 protein beta chain with a conjugated fluorescent tag (Complement C4 polyclonal antibody, AbBy Fluor 647 Conjugated, Bioss, Woburn, MA) for 30 minutes at room temperature with rotation in the dark. The cells were washed with FACS buffer (1% bovine serum albumin and 0.1% sodium azide in phosphate-buffered saline, pH 7.2) and centrifuged (500 × g for 5 minutes at room temperature). Flow cytometry was performed using a Symphony Instrument at the Stanford Shared FACS Facility with the following detectors: B780, B515, V710, V610, U379, R780, R670, Y586, and Y670. All five lasers (488 nm, 405 nm, 355 nm, 638 nm and 561 nm) were used. UltraComp eBeads Plus Compensation Beads (Invitrogen) were prepared with single-antibody controls and used as single-color compensation controls. A randomly chosen donor blood sample was used for the Fluorescence Minus One (FMO) controls and isotype control (Rabbit IgG Isotype Control, AbBy Fluor 647 Conjugated, Bioss, Woburn, MA, USA). HepG2, Jurkat, and genetically modified U937 cell lines were fixed, permeabilized, and stained with C4 antibody as negative and positive controls (Supplementary Materials–Reagent Validation Section, including Supplementary Figures 1 and 2). Supplementary Figure 3 shows the gating strategy used to identify the major immune cell populations: Neutrophils, Natural Killer (NK) cells, T cells, B cells, and monocyte subsets, specifically CD14^+^CD16^−^ (classical monocytes, (CM)), CD14^−^CD16^+^ (nonclassical monocytes, (NCM)), and CD14^+^CD16^+^ (intermediate monocytes). Gates were determined using FMO controls. The

Median Fluorescence Intensity (MFI) was obtained for the signal from the fluorescent tag conjugated to the C4 antibody for each donor. Descriptive statistics for the donors’ C4 protein MFI associated with each defined major immune cell type were determined.

### II: Unbiased examination of C4 gene expression in immune cell types using Meta-signature

More than 50,000 immune cells have been previously analyzed in a meta-analysis of publicly available data (36). Briefly, effect sizes (Hedge’s g) were compared between samples from an immune cell subset of interest and all other immune cell subsets. The resulting gene expression was reported as a Standardized Mean Difference and the relative gene expression of a gene of interest in different immune cells (36). The Meta-Signature Tool (https://metasignature.stanford.edu/) was used to determine the immune cell types expressing C4A and C4B.

### III: The Clinical Comparison Cohort: samples from controls and individuals with SCZ

Samples were collected from two studies: a *previously published Pilot Study* (24) and an *Expanded Cohort*. We referred to the two cohorts as the *Clinical Comparison Cohort*. The inclusion and exclusion criteria and sample handling for the Pilot Cohort were identical to those described for the Expanded Cohort, unless otherwise specified. Participant height and weight were measured to calculate the body mass index (BMI). Age, biological sex, and antipsychotic medication dosage were reported for each participant.

#### Inclusion and exclusion criteria for the Expanded Cohort

For the Expanded Cohort, participants were recruited from a local academic early psychosis clinic and community. Inclusion criteria for the patient group included SCZ or Schizoaffective Disorder diagnosis within the last 5 years as determined by Structured Clinical Interview for the DSM-V by a trained clinician. Healthy controls who met the SCID criteria for a DSM-V diagnosis or had a Distress Score > 2 on the Prodromal Questionnaire were excluded from the study. The study was approved by the Stanford University Institutional Review Board and registered with Clinicaltrials.gov: NCT05109065.

#### Venous blood collection for the Expanded Cohort

Approximately 50 mL of blood was collected in K2 EDTA blood collection tubes via venipuncture between 0800 and 1200 h from overnight fasting participants and processed on the same day. The samples were identified by sample number, blinding the scientists to the case information until the later stages of analysis. The protocols used in the Pilot Study were identical. The freezer storage time (FST) was determined by counting the number of days between sample collection and experimental analysis.

#### Plasma isolation and analysis

Two milliliters of blood were centrifuged within 30 minutes after collection to isolate the plasma. Plasma was snap frozen in liquid nitrogen and stored at -80°C until measurement. Frozen plasma was allowed to thaw on ice and diluted in sample buffer (ProteinSimple, San Jose, CA) 1:500 at 4°C for measurement. Diluted plasma was exposed to C4 alpha chain antibody (22233, Proteintech, Rosemont, IL, USA) with capillary-system-based western WES Protein Simple (Bio-Techne, San Jose, CA). Plasma was diluted 1:500 and C4 antibody was used at 1:50 dilution. The samples were run in duplicate. Care was taken to balance controls and cases in each WES run (each cartridge held approximately 12 samples, analogous to the more familiar ‘gel’). The measured fluorescence intensity of the C4 alpha chain (∼ 90 kDa) was used to compare the C4 protein abundance in the samples. Purified C4 protein was used as a positive control to identify the alpha chain in each run (C4 protein, Comptech, Tyler, TX, USA). C4 protein abundance was normalized using the Sum Normalization method to reduce between cartridge variability (37,38).

#### Peripheral blood mononuclear cell (PBMC) collection for the Expanded Cohort

Peripheral blood mononuclear cells (PBMC) were isolated from room-temperature whole blood using SepMate PBMC Isolation Tubes (StemCell Technologies, Seattle, WA). PBMCs were stored as live cells in liquid nitrogen in fetal bovine serum (FBS) with 10% dimethyl sulfoxide (DMSO). In a Pilot Study, PBMC samples were isolated using the traditional Buffy Coat Method and stored as live cells in liquid nitrogen in FBS with 10% DMSO (24).

#### Neutrophil clinical cohort: a subset of the Clinical Comparison Cohort

A subset of participant samples was processed for neutrophil isolation (Neutrophil Cohort, Supplementary Table 5a). The entire cohort was not included because of limited sample availability. Neutrophils were isolated from fresh whole blood using the EasySep Direct Neutrophil Isolation Kit (StemCell Technologies, Seattle, WA, USA) and stored as cell pellets at -80°C. Pelleted cells were rinsed 1x with phosphate buffered saline (PBS) and pelleted by centrifugation. Neutrophils were not stored as live cells (as PBMCs) because they are known to spontaneously degranulate during the freeze-thaw process(39).

#### Determination of the C4 genotype using digital droplet polymerase chain reaction

Genomic DNA was isolated using standard techniques. The number of copies of the A, B, short (S) and long (L) forms of the C4 gene was determined as previously described (24,25). Measurement distribution was tested for normalcy using the Shapiro-Wilk Test.

### IV: Quantification of C4 protein associated with distinct immune cells

C4 protein was quantified in PBMCs using flow cytometry. Frozen live PBMC were thawed rapidly in 10% FBS in RPMI 1640 media (Gibco, Thermo Fisher Scientific, Pittsburgh, PA) and then collected by centrifugation (300 xg for 5 min at RT). The cells were resuspended in PBS and counted. Next, the cells were stained with the LIVE/DEAD Fixable Violet Dead Cell Stain Kit (Invitrogen, Waltham, MA, USA) according to the manufacturer’s instructions. After washing with FACS buffer (1% bovine serum albumin and 0.1% sodium azide in PBS), the cells were incubated on ice for 15 min in blocking buffer (5% heat-inactivated AB human sera and 5% normal goat serum in PBS). Cells were then stained for exterior stains CD3, CD14, CD16, CD19, CD56, CD66b, CD45, and HLA-DR (same as those described above) by incubation for 30 min at RT in the dark. After washing twice with FACS buffer, cells were fixed and permeabilized using the BD Cytofix/Cytoperm Fixation/Permeabilization Kit (BD Biosciences, San Jose, CA, USA). Finally, cells were stained for C4 protein (bs-15186R-BF647, Bioss, Woburn, MA) after antibody optimization by incubation in antibody in Perm/wash buffer. A parallel sample was stained with an isotype control (bs-0295P-BF647, Bioss, Woburn, MA, USA). Similarly, FMO and compensation controls were freshly prepared for each experiment, as described above. Measurement distribution was tested for normalcy using the Shapiro-Wilk Test. The nonparametric Mann-Whitney U test was used to compare the group means between the SCZ and control samples. Spearman’s correlations were used to test the correlation between measured Neutrophil C4 protein and potential confounders such as age, sex, body mass index (BMI), PBMC freezer storage time (PBMC FST) and antipsychotic medication dosage (converted to olanzapine dose equivalents using the Defined Daily Dose (DDD) Method (40)).

#### Neutrophil C4 protein measurement

Neutrophils were isolated as previously described. Pelleted cells were rinsed 1x with phosphate buffered saline (PBS) and pelleted by centrifugation. Cell pellets were resuspended in cold radioimmunoprecipitation assay buffer (RIPA; Millipore) and allowed to sit for 30 min to ensure cell lysis. Then, the lysates were flash-frozen in liquid nitrogen and thawed on ice. After centrifugation (20,000 × g for 10 min at 4C), the lysate protein concentration was measured using a protein quantification assay (Pierce 600 nm Protein Assay Reagent with the addition of the Ionic Detergent Compatibility Reagent, Thermo Fisher, Pittsburgh, PA). Each samples’ lysate was run in duplicate under optimized conditions to measure abundance of C4 protein (0.5 mg/mL lysate concentration, 1:100 C4 antibody dilution) and actin (0.25 mg/mL lysate concentration, 1:250 actin antibody dilution (beta-actin, 8H10B10, Invitrogen)). The measured fluorescent intensity of the C4 a-chain, ∼90 kDa, normalized by the actin loading control (at ∼48 kDa) was used to compare the C4 protein abundance in samples from controls and individuals with SCZ. Samples were further normalized using the Sum Normalization method to reduce between cartridge variability (37,38). Measurement distribution was tested for normalcy using the Shapiro-Wilk Test. The nonparametric Mann-Whitney U test was used to compare the group means between the SCZ and control samples. Spearman’s correlations were used to test the correlation between measured Neutrophil C4 protein and potential confounders such as age, sex, body mass index (BMI), PBMC freezer storage time (FST) and antipsychotic medication dosage (converted to olanzapine dose equivalents using the Defined Daily Dose (DDD) Method (40)).

#### Classical and nonclassical monocyte isolation, immunofluorescent imaging and analysis

A randomly selected subset of participant samples was processed for monocyte isolation (Monocyte Cohort, Supplementary Table 5b). Monocytes were separated from other PBMC cells using the Pan Monocyte Isolation Kit (Miltenyi, Charlestown, MA, USA) according to the manufacturer’s instructions. Then, CM (CD14^+^CD16^−^) were separated from NCM (CD14^−^CD16^+^) using CD14 Microbeads (Miltenyi, Charlestown, MA, USA). The collected cells were then plated on gelatin-fibronectin-coated chamber slides (ibidi, Fitchburg, WI, USA) in RPMI with 10% FBS. Cells were allowed to incubate at 37°C and 5% carbon dioxide for 30 minutes before being fixed with 4% paraformaldehyde in phosphate buffered saline for 10 minutes. Fixed slides were stored at 4°C until they were stained with Hoescht 34580 (Invitrogen) and antibody directed against the C4 protein (22233-1-AP, Proteintech) using standard methods. Stained samples were imaged using a 63x oil-objective on a Zeiss LSM900 confocal microscope (Courtesy of the Erin Gibson Laboratory, Stanford University). All the samples were imaged with the same field size, exposure, and laser intensity. C4 protein fluorescence was measured across whole cells using a defined region of interest (ROI) in Fiji (41) using the mean fluorescent intensity (mean). The intensity measurements were averaged for the cells from each participant sample. The control and SCZ group means for C4 protein fluorescence intensity were compared on an exploratory basis using the Mann-Whitney U-Test.

#### Statistical analysis

The primary outcome was the correlation between Neutrophil and CM-associated C4 protein and the number of C4A gene copies (GCN) in the control and patient groups. All other statistical tests were considered exploratory and thus were not corrected for multiple comparison. Group differences in samples from individuals with SCZ and controls were compared using ANOVA. Immune cell-associated C4 protein and C4 GCN frequencies were not normally distributed (Shapiro–Wilk test); therefore, nonparametric rank-based tests (Spearman’s rho) were used for correlational analyses. Differences between pairs of rho values were tested for significance using the appropriate modification of Fisher’s *Z*-transformation of correlation coefficients (42,43). All analyses were performed using JMP Pro 14.1.0 (SAS Institute, Cary, NC).

## Results

### C4 protein is primarily localized in neutrophils and monocytes

Ten whole blood samples from anonymous donors were obtained from the Stanford Blood Center (demographic information in Supplementary Table 1). Using flow cytometry, we found C4 protein predominantly in neutrophils and all three types of monocytes (Figure 1c-d). The rabbit IgG isotype control signal was approximately ten times lower than the measured C4 protein signal in all immune cell types, except for neutrophils (Figure 1c-d), suggesting that the antibody has increased nonspecific interactions with neutrophils compared to other immune cell types. The negative controls (FMO and Jurkat cell line, Figure 1a and Supplementary Figure 1) are lower in C4 protein MFI than all the immune cell types and isotype controls. The positive control cell line (hepG2, Figure 1a and Supplementary Figure 1) has a higher C4 protein MFI than all the isotype controls except for the neutrophil isotype control.

**Figure 1.**
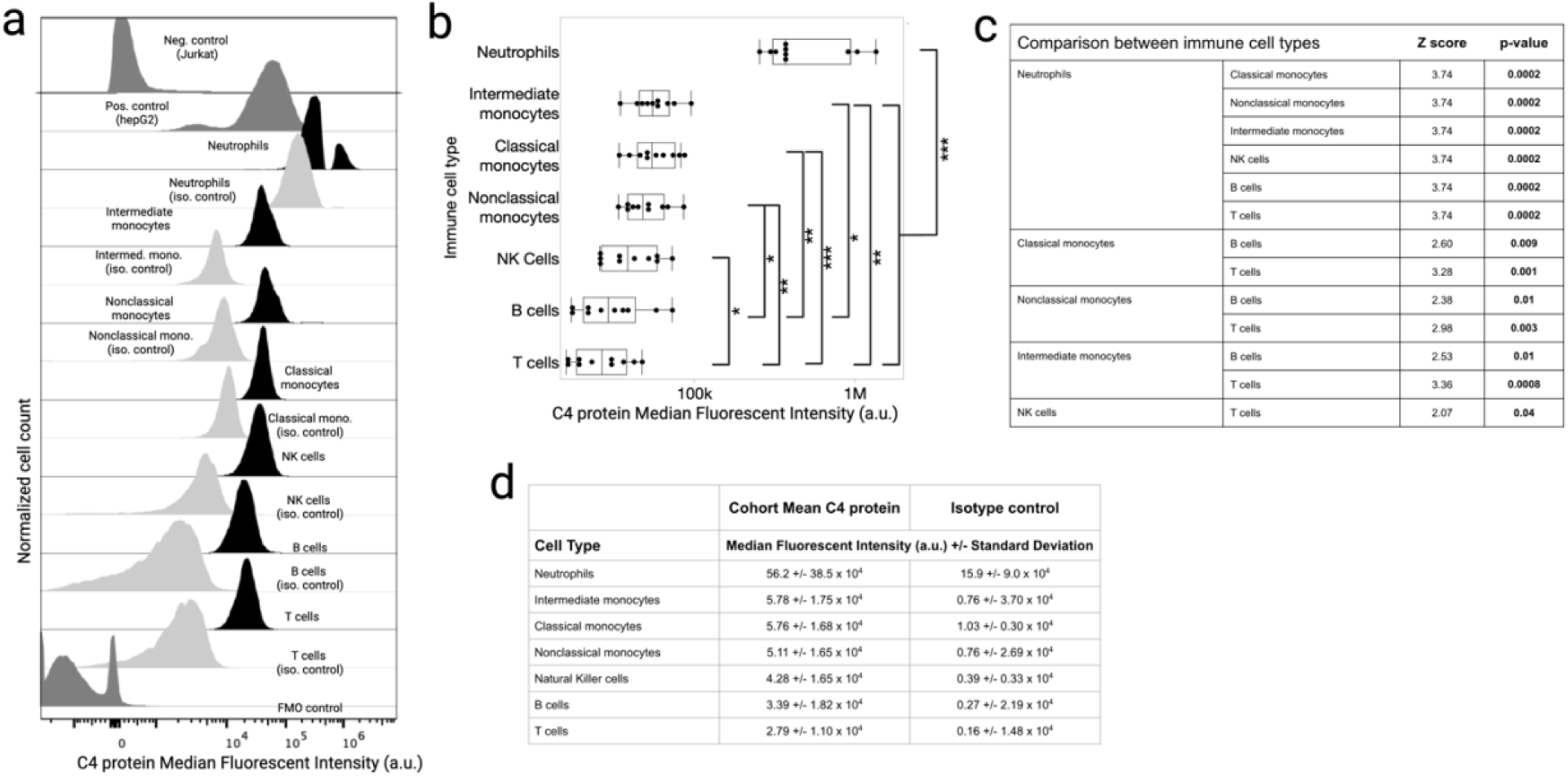
Neutrophils and monocytes express the C4A gene and contain C4 protein. **(a)** Flow cytometry of fresh whole-blood samples from anonymous donors. C4 protein was quantified (Median Fluorescent Intensity) in major immune cell types using flow cytometry. Jurkat cell line is the negative control and hepG2 is the positive control cell line (top 2 panels in dark gray). Rabbit IgG isotype controls for each major immune cell type are shown in the panel below the C4 protein quantification in that immune cell type (light gray). The bottom panel shows FMO control (dark gray). **(b)** Median Fluorescence Intensity (MFI) in major immune cell types in ten healthy donors from the Stanford Blood Center, as measured by flow cytometry. Group differences between immune cell types were tested using the nonparametric Mann-Whitney U Test and significance was tested using the false discovery rate (FDR). * = p <0.05, ** = p <0.01, *** = p <0.001. **(c)** Comparisons that survived threshold for multiple comparisons are shown. A comprehensive list of comparisons is provided in Supplementary Table 3. **(f)** Average of Median Fluorescence Intensity (MFI) of C4 protein signal in major immune cell types in ten healthy donors from the Stanford Blood Center. An isotype control from the rabbit IgG isotype control for each immune cell type was also provided.

Nevertheless, the amount of C4 protein associated with neutrophils was higher than C4 protein associated with the other immune cells tested (Figure 1c-d, Statistics shown in Supplementary Table 3, p = 0.0002). Likewise, C4 protein associated with all three identified monocyte subtypes (CM, NCM, and intermediate) was higher than that in T cells (p = 0.001 for CM, p=0.003 for NCM, and p = 0.0008 for intermediate monocytes) and B cells (p = 0.009 in CM, p=0.01 in NCM, and p = 0.01 in intermediate monocytes) (Figure 1c-d, Supplementary Table 3). Lastly, natural killer (NK) cell MFI values were higher than those of T cells (p = 0.04).

Limited demographic information was available for the donors (Supplementary Table 1a). No significant correlations were observed between C4 protein MFI in the major immune cell types and age (Supplementary Table 1b).

### C4A and C4B genes are expressed in different cell types

C4A and C4B gene expression varied among different immune cell types (Supplementary Figure 3a-b). The C4A gene is expressed approximately twice as much in intermediate (CD14^+^CD16^+^) monocytes, neutrophils, specialized memory T cells, and dendritic cells as compared to other immune cells (Supplementary Figure 3a). In contrast, the C4B gene was expressed at approximately three times the amount in T and B cells compared to other immune cells (Supplementary Figure 3b). CM (CD14^+^) express less C4A and C4B genes compared to other immune cell types; however, they express twice the amount of the C4A gene compared to the C4B gene.

### Demographic and clinical characteristics of SCZ and control participants

A total of 38 controls and 25 individuals with SCZ agreed to enroll in the study. Seven volunteers in the control group were ineligible after the SCID-V assessment due to a current psychiatric disorder. Three individuals with SCZ were ineligible after SCID-V assessment due to meeting the criteria for a substance abuse disorder or not meeting the criteria for SCZ or schizoaffective disorder. One control and two patients withdrew before completing the study. Thus, 30 healthy controls and 20 cases completed clinical evaluation and blood sampling (Table 1 and Supplementary Table 4). One control subject had to be excluded because of insufficient sample quantity. Additionally, nine controls and 10 individuals with SCZ were included in our pilot study (24). A total of 38 control and 30 individuals with SCZ were included in the analysis. The two groups did not differ significantly with respect to age, sex, or ethnicity.

**Table 1:**
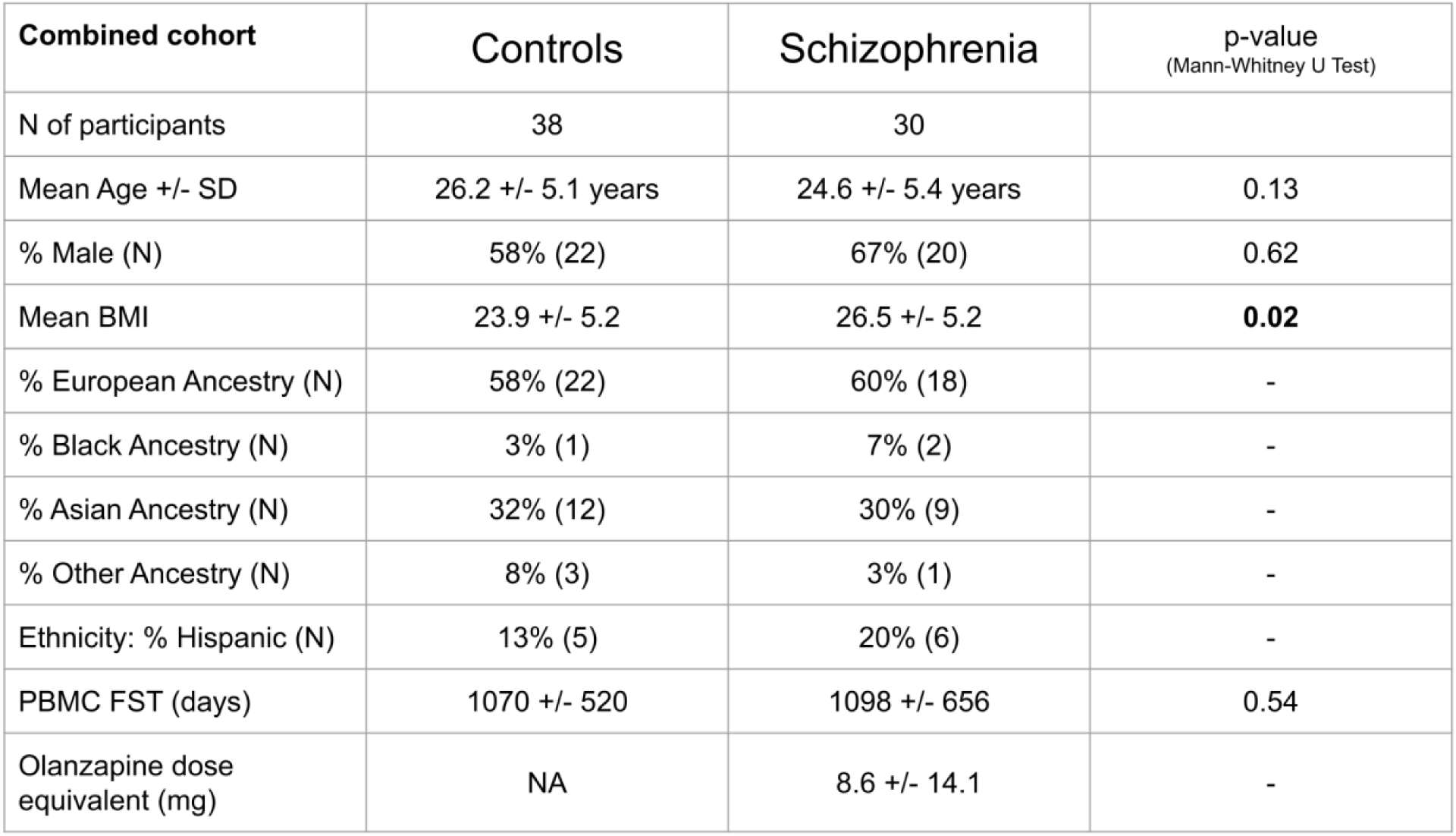
Demographics of the study participants in the Clinical Comparison Cohort. Summary statistics for Age, Sex (percentage and number of male participants), Body Mass Index (BMI), Plasma Freezer Storage Time, PBMC Freezer Storage Time and Ethnicity are provided for control and individuals with SCZ and Control groups. The distributions were compared using the Mann-Whitney U test. N = Number, SD = Standard Deviation, BMI = Body Mass Index, PBMC = Peripheral Blood Mononuclear Cells, FST = Freezer Storage Time. ^+^Clinical Comparison Cohort which consists of the Pilot and Expanded Cohorts together.

Individuals with SCZ had a higher body mass index (BMI) than controls (23.9 ± 5.2, 26.5 ± 5.2, p = 0.02; Table 1), as is expected for this population (44). None of the control participants were taking any medications when enrolled in the study. All individuals with SCZ were on medication: 97.2% were taking antipsychotic medication, 34.3% were taking more than one antipsychotic medication, 37% were taking antidepressants, 28.6% were taking medications for anxiety, and 8.6% were taking mood stabilizers. Additionally, 14.3% were taking benztropine and 2.9% were taking samidorphan.

Subsets of the Clinical Comparison Cohort were used for neutrophil and monocyte experiments, based on sample availability. The demographics of the study participants comprising the Neutrophil and Monocyte Subsets are provided in Supplementary Table 5. They did not differ from the Clinical Comparison Cohort in terms of the metrics evaluated (age, sex, BMI, antipsychotic medication, FST, or ethnicity).

### Neutrophil C4 protein content correlate with C4A gene copy number in individuals with SCZ

We investigated whether C4 protein within major immune cell types (neutrophils and monocytes) correlated with the number of C4A gene copies in SCZ and/or controls. To test this, we determined the number of C4A genes using ddPCR and C4 protein associated with neutrophils using western blotting, and in CM from live PBMCs using flow cytometry (Figure 2 and Figure 3). Correlations were tested using CM only, since the C4 protein associated with immune cells was similar in different monocyte subsets (Figure 1c-d), and CM are the dominant monocyte subtype in blood, constituting approximately 80% of total monocytes. In our cohort, we found a higher number of total C4, C4A, and C4AL gene copies in the SCZ group than in the control group (Figure 2b). We found a moderate positive correlation between the number of C4A gene copies and C4 protein *only* in Neutrophils and *only* in the SCZ group (r = 0.64, p = 0.01, Figure 2c). We observe a trend towards a difference in correlations between SCZ samples and controls (*z*[difference] = −1.68, *p* = 0.09). We did not find a correlation between C4 protein in plasma or CM and the number of C4A gene copies in either the control or SCZ groups (Figure 2d).

**Figure 2.**
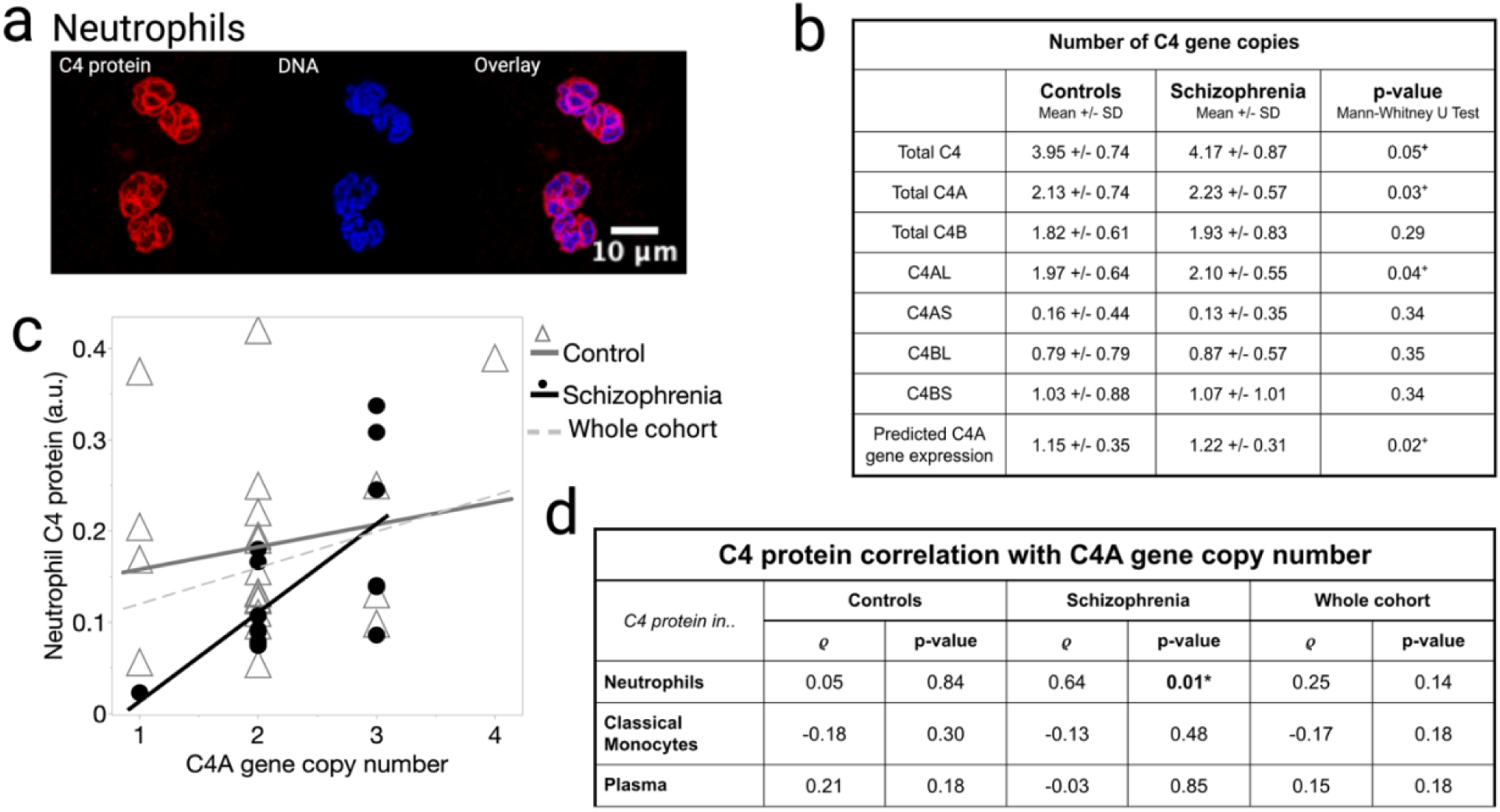
C4 protein in neutrophils is positively correlated with the number of C4A gene copies in SCZ. **(a)** Isolated neutrophils from a healthy donor labeled for C4 protein (show in red, labeled with anti-C4 antibody (Proteintech 22233)) and DNA (shown in blue, stained with DAPI). The overlay shows that the C4 protein is at the rim of the nuclear DNA. **(b)** The Number of C4 gene copies for the various forms of the C4 gene (AS, AL, BS, and BL) was determined using ddPCR. Descriptive statistics for the control and SCZ groups in the Clinical Comparison Cohort are also provided. **(c)** Spearman correlation between measured Neutrophil C4 protein and the number of C4A gene copies is provided for the control, SCZ, and whole cohorts. The correlation between C4 protein associated with Neutrophils and the number of C4A gene copies was found to be statistically significant (r = 0.64, p = 0.01). The difference in correlations in individuals with SCZ and in controls trended towards significance based on the modified Fisher’s *Z*-transformation test for rho (*z*[difference] = −1.68, *p* = 0.09). **(d)** Table showing the Spearman’s correlation of the number of C4A gene copies and the measured C4 protein associated with Neutrophils and CM in the SCZ, control, and whole (combined SCZ and control) cohorts.

**Figure 3.**
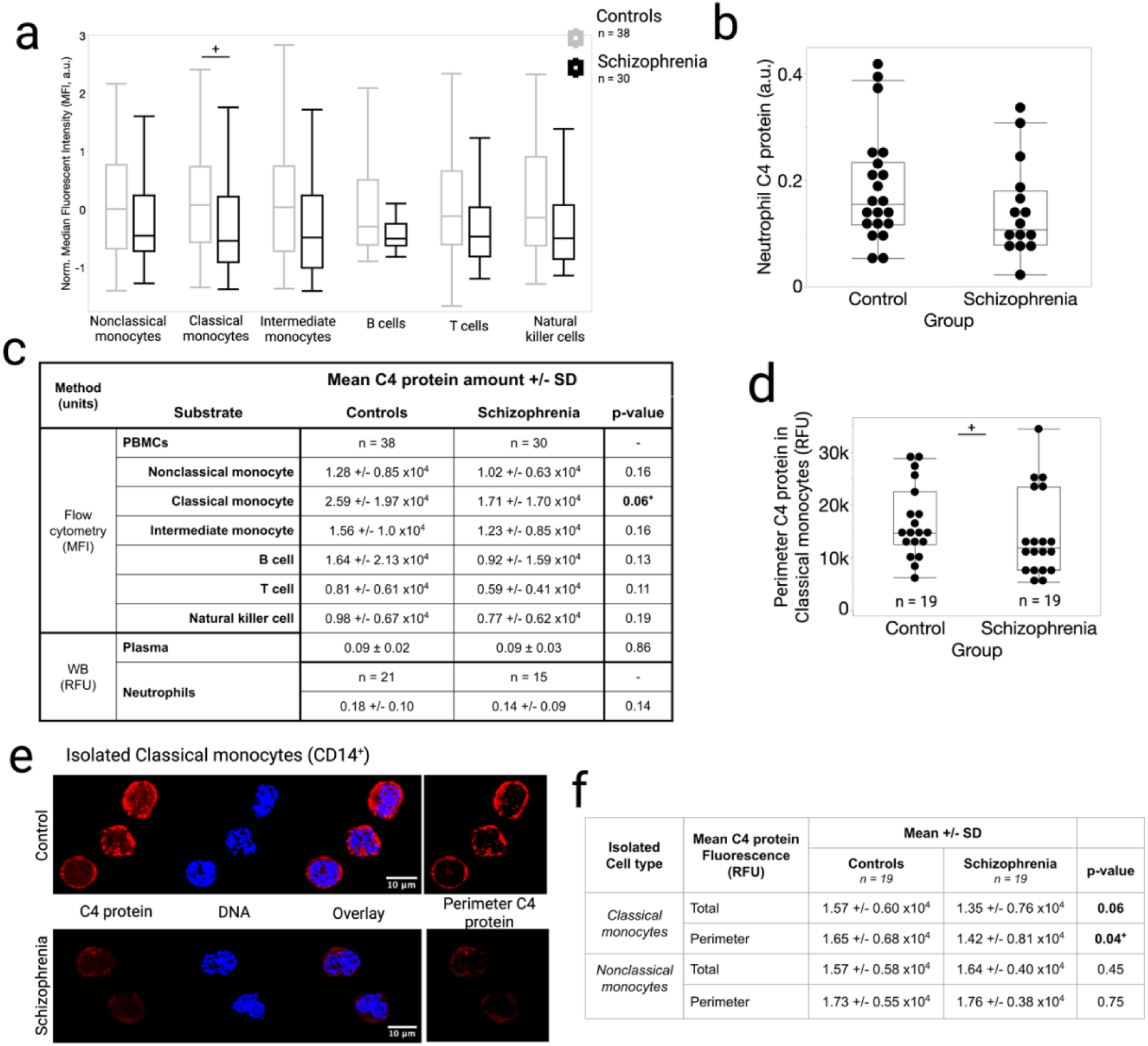
C4 protein is localized to the periphery of classical monocytes and is decreased in individuals with SCZ compared to Controls. **(a)** Flow cytometry was used to quantify C4 protein in major immune cell subtypes. **(b)** C4 protein was measured by immunoblotting against C4 protein (22233 Proteintech, Rosemont IL) using WES (capillary-based western blotting) in isolated neutrophils sampled from 15 SCZ and 21 control individuals. Samples were normalized using cellular actin (8H10D10, Invitrogen, Waltham, MA, USA). **(c)** Table showing the Descriptive statistics of measured immune cell-associated C4 protein in samples from individuals with SCZ compared to controls using different methods. Group comparisons of C4 protein for each major immune cell type (and plasma) were performed using one-way ANOVA. **(d-f)** CM and NCM were isolated from frozen live PBMC samples. Isolated cells were incubated, fixed, and stained for C4 protein (antibody directed against C4 protein, Proteintech, 22233-1-AP). C4 protein is localized throughout monocyte cells but is preferentially localized at the periphery. Representative images of CMs stained with Hoeschet stain for DNA and fluorescently labeled antibody against C4 protein in a sample from a control participant and a sample from a participant with SCZ. **(e)** Quantification of C4 protein throughout each cell was performed by measuring the Mean Intensity (Mean) from immunofluorescent images of C4 protein using Fiji. A mask was created from the nuclear stain and used to subtract the central C4 protein fluorescence to determine the Peripheral C4 protein Mean Fluorescence. **(f)** Table showing the exploratory descriptive statistics of CM and NCM C4 protein in samples from individuals with SCZ compared to controls. RFU = Relative Fluorescent Unit.

We examined the correlation between Neutrophil C4 protein amount and potential confounders (Supplementary Figure 5). In the control group, we found a moderate positive correlation between Neutrophil C4 protein and both age (r = 0.46, p = 0.04, Supplementary Figure 5a-b) and BMI (r = 0.40, p = 0.07, Supplementary Figure 5a,c).

### C4 protein may be depleted in CM from individuals with Schizophrenia

We conducted an exploratory analysis to compare whether the amount of C4 protein associated with specific immune cell types (neutrophils, NCM, CM, intermediate monocytes, B cells, T cells, and natural killer cells) and plasma differed between SCZ samples and controls. C4 protein levels in plasma and neutrophils were compared using western blotting. C4 protein levels in other immune cell subtypes were compared using flow cytometry from previously frozen live PBMCs. We observed decreased C4 protein across *all* immune cells in SCZ samples compared to controls (Figure 3a-c). The largest difference was in C4 protein associated with CM (MFI = 2.59 +/- 1.97 ×10^4^ in controls and 1.71 +/- 1.70 ×10^4^ in SCZ samples; p = 0.06).

To further explore whether C4 protein associated with CM is altered in SCZ samples, we isolated, plated, and stained CM (with NCM as a control) for C4 protein. C4 protein immunofluorescence also allowed us to determine the localization of C4 protein in CM and NCM. We observed that C4 protein was found at the perimeter of the cell and throughout the cytoplasm (Figure 3e). Next, we compared the group means of total C4 protein (throughout the whole cell) and C4 protein at the perimeter, measured by the Mean Relative Fluorescent Intensity (RFU). We found significantly lower peripheral C4 protein RFU in SCZ CM (1.42 +/- 0.81 × 10^4^ in SCZ samples vs. 1.65 +/- 0.68 × 10^4^ in controls, p = 0.04). Likewise, we found a trend towards lower total (whole cell) C4 protein RFU in SCZ CM compared to controls (1.42 +/- 0.81 × 10^4^ in SCZ samples vs. 1.65 +/- 0.68 × 10^4^ in controls, p = 0.06). Conversely, we did not find a difference between SCZ and control samples in NCM in either total or peripheral C4 protein RFU.

We examined the correlations between CM C4 protein and potential confounders (Supplementary Figure 6). In CM C4 protein using flow cytometry, we find a small inverse correlation between CM C4 protein and FST (controls: r = -0.26, p = 0.12, SCZ: r = -0.30, p = 0.10, and whole cohort: r = -0.32, p = 0.01, Supplementary Figure 6a-b). Conversely, we find a moderate positive correlation between CM C4 protein and FST in the SCZ and whole cohort (SCZ: r = 0.46, p = 0.05, and whole cohort: r = 0.32, p = 0.05, Supplementary Figure 6c-d).

## Discussion

The absence of treatments for schizophrenia underscores the need to better understand its underlying pathophysiology. Peripheral immune mechanisms represent a promising area for study. Treatment targets in the periphery are attractive because pharmaceutical agents do not need to cross the blood-brain barrier and still have therapeutic effects on the brain (19,20). Innate immune mechanisms are a promising lead since they follow the disease course; particularly around C4 protein activation (C4-ana) (6,11,15,30). In support of our overarching hypothesis, the presence of C4 protein in neutrophils and monocytes provides a non-plasma source of potential C4 protein activation in SCZ. We demonstrated that C4 protein is expressed by and present in neutrophils and monocytes (Figure 1, Supplementary Figure 4). Furthermore, when we compared the correlation between neutrophil C4 protein and the number of C4A gene copies, we found a moderate positive correlation *only* in the SCZ group (Figure 2). We did not find a correlation between C4 protein and the number of C4A gene copies for either group in CM or plasma. Likewise, we did not find a correlation between C4 protein and the number of C4B gene copies in neutrophils, CM, or plasma in either group. These data suggest that SCZ neutrophils actively express the C4A gene and protein. This interpretation is consistent with meta-analyses of cytokines in CHR, FEP and chronic SCZ which are elevated in SCZ samples and inducers of C4A gene expression, like interferon-g and IL-6 (3,45). Because C4 protein is expressed but does not accumulate in SCZ neutrophils, it is either consumed in a biological pathway or excreted. We previously found a positive correlation between C4-ana (C4 protein activation product) and the number of C4A gene copies (24). Thus, we favor the interpretation that C4 protein is consumed in a biological pathway. If C4 protein was excreted, we would expect to find higher amounts of C4 protein in plasma in SCZ samples because neutrophils make up about half (∼40-60%) of all white blood cells in the peripheral circulation. We did not find higher amounts of C4 protein in plasma from SCZ samples compared to controls, which is consistent with previous studies (Figure 2).

In our exploratory analyses, we observed a decrease in C4 protein in SCZ CM, particularly at the perimeter (Figure 3). While not statistically significant, this trend was consistently observed using two different methods of measuring C4 protein in CM and two different antibodies that target the C4 protein (Figure 3a,d-f). The decrease in C4 protein was more pronounced at the cell perimeter (Figure 3d-f).

However, the interpretation of decreased C4 protein in SCZ CM is less clear. Similar to neutrophils, CM are also known to be in an activated state in SCZ samples (46). However, we did not observe a correlation between the amount of C4 protein in CM and the number of C4A gene copies, suggesting that C4 protein is not actively expressed in CM. Thus, we hypothesized that C4 protein is consumed as part of CM activation in SCZ samples.

The detection of C4 protein inside neutrophils and monocytes is a novel basic scientific finding. Our data support the existence of a C4 complosome, an intracellular C4 protein (36). Neutrophils and monocytes express the C4A gene and protein (Supplementary Figure 4). However, the function of C4 protein in these immune cell types is unknown. We hypothesize that C4 protein is part of the biological pathways involved in neutrophil and monocyte activation because we see evidence of either consumption (in monocytes) and/or active transcription (in neutrophils) in SCZ samples, which have been shown to be in an activated state. Further research on the fundamental mechanisms underlying the function of neutrophils and monocytes is warranted.

This study has several limitations. First, we used fresh whole blood, which required us to use blood donated on the day we performed the experiment (Figure 1). Blood donors may have chronic conditions and are allowed to be on medication. The number of volunteers was small. In contrast, the C4 protein measured in the blood samples from biobank volunteers was largely in agreement with the gene expression data obtained from a much larger population (>800 samples, Figure 1a). Another limitation is the high affinity of the isotype control antibody to neutrophils, raising the concern that the amount of C4 protein per neutrophil may be inflated owing to nonspecific binding of the antibody (Figure 1c, f). The high binding affinity of the isotype control appeared to be isolated to neutrophils, as evidenced by the 10x lower MFI signal of the isotype control compared to the other immune cells (Figure 1c,f). The negative control cell line (Jurkat) had a near-zero C4 protein MFI signal (Figure 1b). Taken together, we can reasonably conclude that neutrophils express and contain high amounts of C4 protein, although the exact quantification of C4 protein per neutrophil relative to other cell types may be imprecise.

In our Clinical Comparison Cohort (Table 1), the interpretation of our study was limited by potential confounders such as medication use, FST, and high BMI in participants with SCZ. While we did not detect an association between either Neutrophil or CM C4 protein and medication use, this study should be repeated in samples from medication-naïve FEP that are known to progress to SCZ to determine whether this association remains without the potential confounding of medication exposure. The associations that we detected between Neutrophil C4 protein and age and BMI were limited to the control group (associations between immune system changes in obesity and aging are well known (45,46)), supporting the claim that the changes we found in Neutrophil C4 protein in SCZ samples are more reflective of the disease state of SCZ (Supplementary Figure 4). In examining the potential associations between CM C4 protein and common confounders, we found a small negative correlation between CM C4 protein measured by flow cytometry and FST (Supplementary Figure 4). Monocytes are known to be affected by freeze-thaw (47). The same small effect was observed in both the control and SCZ groups. In the CM C4 protein measured by immunofluorescence, we observed a small negative correlation between CM C4 protein and FST in the SCZ group (Supplementary Figure 5). The fact that the directionality is opposite to that of CM C4 protein and FST measured by flow cytometry suggests that the association arises from the methods used to process the samples. CM were isolated prior to immunofluorescence, possibly selecting for a healthier subset of SCZ CM. Selection of a healthier population of CM from SCZ samples suggests that the loss of C4 protein associated with CM may be more pronounced than what we are able to measure in this study.

The finding of C4 protein in neutrophils (Figure 1) and the positive association with the number of C4A gene copies (Figure 2) support the hypothesis that the C4 protein in these cells could be the source of complement activation which is consistently observed in SCZ samples. These findings integrate several previously separate aspects of innate immunity and observations in SCZ that follow disease-related features: the risk association of SCZ GWAS loci (number of copies of the C4A gene), evidence of complement activation in the plasma around the C4 protein, and the primary cells of the innate immune response (neutrophils and monocytes). Similarly, it is well known that the most effective current medication for the treatment of SCZ, clozapine, inhibits neutrophils (48,49). Taken together, the convergence of these disparate factors strongly suggests that we may be honing a key pathophysiological mechanism of SCZ pathophysiology in the peripheral circulation, which we hope will lead to novel, accessible disease-altering therapeutics.

## Supporting information

All supp materials

## Funding sources

Stanford University Department of Psychiatry and Behavioral Sciences Innovator Award (to AK)

Stanford University Department of Medicine Translational Research and Applied Medicine Award (to AK)

VISN21 Veterans Administration Early Career Award Program (to AK)

K08MH132042 (to AK)

Data was collected on an instrument in the Shared FACS Facility obtained using NIH S10 Shared Instrument Grant 1S100D026831-01.

## Acknowledgements

Erin Gibson laboratory at Stanford University Department of Psychiatry and Behavioral Sciences for the generous sharing of confocal microscope for imaging in this project.

Figures Created in BioRender. Kalinowski, A. (2025) https://BioRender.com/p1o76kc; https://BioRender.com/sj7bs41

## References

1. Estes ML, McAllister AK. Maternal immune activation: Implications for neuropsychiatric disorders. Science. 2016;353(6301):772–7.

2. Khandaker GM, Cousins L, Deakin J, Lennox BR, Yolken R, Jones PB. Inflammation and immunity in schizophrenia: Implications for pathophysiology and treatment. Lancet Psychiatry. 2015;2(3):258–70.

3. Upthegrove R, Khandaker GM. Cytokines, Oxidative Stress and Cellular Markers of Inflammation in Schizophrenia. In: Khandaker GM, Meyer U, Jones PB, editors. Neuroinflammation and Schizophrenia [Internet]. Cham: Springer International Publishing; 2019 [cited 2024 Aug 29]. p. 49–66. (Current Topics in Behavioral Neurosciences; vol. 44). Available from: https://link.springer.com/10.1007/7854_2018_88

4. Karlsson H, Dalman C. Epidemiological Studies of Prenatal and Childhood Infection and Schizophrenia. Curr Top Behav Neurosci. 2020;44:35–47.

5. Orlovska-Waast S, Köhler-Forsberg O, Brix SW, Nordentoft M, Kondziella D, Krogh J, et al. Cerebrospinal fluid markers of inflammation and infections in schizophrenia and affective disorders: a systematic review and meta-analysis. Mol Psychiatry. 2019;24(6):869–87.

6. Dudeck L, Nussbaumer M, Nickl-Jockschat T, Guest PC, Dobrowolny H, Meyer-Lotz G, et al. Differences in Blood Leukocyte Subpopulations in Schizophrenia: A Systematic Review and Meta-Analysis. JAMA Psychiatry [Internet]. 2025 Mar 5 [cited 2025 Mar 6]; Available from: https://jamanetwork.com/journals/jamapsychiatry/fullarticle/2830862

7. Gallego JA, Blanco EA, Husain-Krautter S, Fagen EM, Moreno-Merino P, Ojo-Jiménez JAD, et al. Cytokines in cerebrospinal fluid of patients with schizophrenia spectrum disorders: New data and an updated meta-analysis. Schizophr Res. 2018;202:64–71.

8. Halstead S, Siskind D, Amft M, Wagner E, Yakimov V, Liu ZSJ, et al. Alteration patterns of peripheral concentrations of cytokines and associated inflammatory proteins in acute and chronic stages of schizophrenia: a systematic review and network meta-analysis. Lancet Psychiatry. 2023 Apr;10(4):260–71.

9. Mondelli V, Forti MD, Morgan BP, Murray RM, Pariante CM, Dazzan P. Baseline high levels of complement component 4 predict worse clinical outcome at 1-year follow-up in first-episode psychosis. Brain Behav Immun. 2020;(December 2019):0–1.

10. Mongan D, Sabherwal S, Susai SR, Föcking M, Cannon M, Cotter DR. Peripheral complement proteins in schizophrenia: A systematic review and meta-analysis of serological studies. Schizophr Res. 2020;222:58–72.

11. Byrne JF, Healy C, Föcking M, Heurich M, Susai SR, Mongan D, et al. Plasma complement and coagulation proteins as prognostic factors of negative symptoms: An analysis of the NAPLS 2 and 3 studies. Brain Behav Immun. 2024 Jul;119:188–96.

12. English JA, Lopez LM, O’Gorman A, Föcking M, Hryniewiecka M, Scaife C, et al. Blood-Based Protein Changes in Childhood Are Associated With Increased Risk for Later Psychotic Disorder: Evidence From a Nested Case-Control Study of the ALSPAC Longitudinal Birth Cohort. Schizophr Bull. 2018;44(2):297–306.

13. Williams JA, Burgess S, Suckling J, Lalousis PA, Batool F, Griffiths SL, et al. Inflammation and Brain Structure in Schizophrenia and Other Neuropsychiatric Disorders: A Mendelian Randomization Study. JAMA Psychiatry. 2022 May 1;79(5):498.

14. Ji E, Boerrigter D, Cai HQ, Lloyd D, Bruggemann J, O’Donnell M, et al. Peripheral complement is increased in schizophrenia and inversely related to cortical thickness. Brain Behav Immun [Internet]. 2021 Nov; Available from: http://www.ncbi.nlm.nih.gov/pubmed/34808287

15. Susai SR, Föcking M, Mongan D, Heurich M, Coutts F, Egerton A, et al. Association of Complement and Coagulation Pathway Proteins With Treatment Response in First-Episode Psychosis: A Longitudinal Analysis of the OPTiMiSE Clinical Trial. Schizophr Bull. 2023 Jul 4;49(4):893–902.

16. Susai SR, Mongan D, Healy C, Cannon M, Cagney G, Wynne K, et al. Machine learning based prediction and the influence of complement – Coagulation pathway proteins on clinical outcome: Results from the NEURAPRO trial. Brain Behav Immun. 2022 Jul;103:50–60.

17. Wheeler MA, Quintana FJ. The neuroimmune connectome in health and disease. Nature. 2025 Feb 13;638(8050):333–42.

18. Cathomas F, Holt LM, Parise EM, Liu J, Murrough JW, Casaccia P, et al. Beyond the neuron: Role of non-neuronal cells in stress disorders. Neuron. 2022 Apr 6;110(7):1116–38.

19. Middeldorp J, Lehallier B, Villeda SA, Miedema SSM, Evans E, Czirr E, et al. Preclinical Assessment of Young Blood Plasma for Alzheimer Disease. JAMA Neurol. 2016 Nov 1;73(11):1325.

20. Sha SJ, Deutsch GK, Tian L, Richardson K, Coburn M, Gaudioso JL, et al. Safety, Tolerability, and Feasibility of Young Plasma Infusion in the Plasma for Alzheimer Symptom Amelioration Study: A Randomized Clinical Trial. JAMA Neurol. 2019 Jan 1;76(1):35.

21. Weickert TW, Jacomb I, Lenroot R, Lappin J, Weinberg D, Brooks WS, et al. Adjunctive canakinumab reduces peripheral inflammation markers and improves positive symptoms in people with schizophrenia and inflammation: A randomized control trial. Brain Behav Immun. 2024 Jan;115:191–200.

22. Girgis RR, Ciarleglio A, Choo T, Haynes G, Bathon JM, Cremers S, et al. A Randomized, Double-Blind, Placebo-Controlled Clinical Trial of Tocilizumab, An Interleukin-6 Receptor Antibody, For Residual Symptoms in Schizophrenia. Neuropsychopharmacology. 2018 May;43(6):1317–23.

23. Muller N, Myint AM, J. Schwarz M. Immunological Treatment Options for Schizophrenia. Curr Pharm Biotechnol. 2012 May 1;13(8):1606–13.

24. Kalinowski A, Liliental J, Anker LA, Linkovski O, Culbertson C, Hall JN, et al. Increased activation product of complement 4 protein in plasma of individuals with schizophrenia. Transl Psychiatry. 2021;11(1):486.

25. Sekar A, Bialas AR, Rivera HD, Davis A, Hammond TR, Kamitaki N, et al. Schizophrenia risk from complex variation of complement component 4. Nature. 2016;530(7589):177–83.

26. Yilmaz M, Yalcin E, Presumey J, Aw E, Ma M, Whelan CW, et al. Overexpression of schizophrenia susceptibility factor human complement C4A promotes excessive synaptic loss and behavioral changes in mice. Nat Neurosci [Internet]. 2020; Available from: 10.1038/s41593-020-00763-8

27. Sellgren CM, Gracias J, Watmuff B, Biag JD, Thanos JM, Whittredge PB, et al. Increased synapse elimination by microglia in schizophrenia patient-derived models of synaptic pruning. Nat Neurosci. 2019;22(3):374–85.

28. Gracias J, Orhan F, Hörbeck E, Holmén-Larsson J, Khanlarkani N, Malwade S, et al. Cerebrospinal fluid concentration of complement component 4A is increased in first episode schizophrenia. Nat Commun. 2022 Nov;13(1):6427.

29. Koskuvi M, Malwade S, Gracias Lekander J, Hörbeck E, Bruno S, Holmen Larsson J, et al. Lower complement C1q levels in first-episode psychosis and in schizophrenia. Brain Behav Immun. 2024 Mar;117:313–9.

30. Mongan D, Föcking M, Healy C, Susai SR, Heurich M, Wynne K, et al. Development of Proteomic Prediction Models for Transition to Psychotic Disorder in the Clinical High-Risk State and Psychotic Experiences in Adolescence. JAMA Psychiatry. 2020;

31. Mayilyan KR, Weinberger DR, Sim RB. The complement system in schizophrenia. Drug News Perspect. 2008;21(4):200–10.

32. Woo JJ, Pouget JG, Zai CC, Kennedy JL. The complement system in schizophrenia: where are we now and what’s next? Mol Psychiatry. 2020;25(1):114–30.

33. Alper CA, Johnson AM, Birtch AG, Moore FD. Human C′3: Evidence for the Liver as the Primary Site of Synthesis. Science. 1969 Jan 17;163(3864):286–8.

34. Morgan BP, Gasque P. Extrahepatic complement biosynthesis: where, when and why? Clin Exp Immunol. 2003 Nov 14;107(1):1–7.

35. West EE, Kemper C. Complosome — the intracellular complement system. Nat Rev Nephrol. 2023 Jul;19(7):426–39.

36. Vallania F, Zisman L, Macaubas C, Hung SC, Rajasekaran N, Mason S, et al. Multicohort Analysis Identifies Monocyte Gene Signatures to Accurately Monitor Subset-Specific Changes in Human Diseases. Front Immunol. 2021;12:659255.

37. Degasperi A, Birtwistle MR, Volinsky N, Rauch J, Kolch W, Kholodenko BN. Evaluating Strategies to Normalise Biological Replicates of Western Blot Data. Vrana KE, editor. PLoS ONE. 2014 Jan 27;9(1):e87293.

38. Rees PA, Lowy RJ. Optimizing reduction of western blotting analytical variations: Use of replicate test samples, multiple normalization methods, and sample loading positions. Anal Biochem. 2023 Aug;674:115198.

39. Takahashi T, Inada S, Pommier CG, O’Shea JJ, Brown EJ. Osmotic stress and the freeze-thaw cycle cause shedding of Fc and C3b receptors by human polymorphonuclear leukocytes. J Immunol Baltim Md 1950. 1985 Jun;134(6):4062–8.

40. Leucht S, Samara M, Heres S, Davis JM. Dose Equivalents for Antipsychotic Drugs: The DDD Method: Table 1. Schizophr Bull. 2016 Jul;42(suppl 1):S90–4.

41. Schindelin J, Arganda-Carreras I, Frise E, Kaynig V, Longair M, Pietzsch T, et al. Fiji: an open-source platform for biological-image analysis. Nat Methods. 2012 Jul;9(7):676–82.

42. Sheskin D. Handbook of parametric and nonparametric statistical procedures. Third. Chapman & Hall/CRC; 2004.

43. Zarr J. Biostatistical Analysis. Prentice H. 1999.

44. Pillinger T, McCutcheon RA, Vano L, Mizuno Y, Arumuham A, Hindley G, et al. Comparative effects of 18 antipsychotics on metabolic function in patients with schizophrenia, predictors of metabolic dysregulation, and association with psychopathology: a systematic review and network meta-analysis. Lancet Psychiatry. 2020 Jan;7(1):64–77.

45. Upthegrove R, Manzanares-Teson N, Barnes NM, Goldsmith DR, Rapaport MH, Miller BJ, et al. Cytokine alterations in first-episode schizophrenia and bipolar disorder: Relationships to brain structure and symptoms. Schizophr Bull. 2018;15(1):1696–709.

46. Kübler R, Ormel PR, Sommer IEC, Kahn RS, De Witte LD. Gene expression profiling of monocytes in recent-onset schizophrenia. Brain Behav Immun. 2023 Jul;111:334–42.

47. Haider P, Hoberstorfer T, Salzmann M, Fischer MB, Speidl WS, Wojta J, et al. Quantitative and Functional Assessment of the Influence of Routinely Used Cryopreservation Media on Mononuclear Leukocytes for Medical Research. Int J Mol Sci. 2022 Feb 7;23(3):1881.

48. Buchanan RW. Clozapine: Efficacy and Safety. Schizophr Bull. 1995 Jan 1;21(4):579–91.

49. Capannolo M, Fasciani I, Romeo S, Aloisi G, Rossi M, Bellio P, et al. The atypical antipsychotic clozapine selectively inhibits interleukin 8 (IL-8)-induced neutrophil chemotaxis. Eur Neuropsychopharmacol. 2015 Mar;25(3):413–24.

