## Supplementary material for "Peripheral Complement C4 Protein in Schizophrenia: Association with Gene Copy Number and Immune Cell Subtypes": All supp materials

### Supplementary Materials

#### 1. Reagent Validation

##### A. Identifying positive and negative control cell lines

To identify positive and negative controls for our experiments, we tested C4A/B mRNA expression and C4 protein abundance in common cell lines available in our laboratories.

**Methods:**

*Measurement of C4A and C4B gene expression* To determine mRNA expression, we grew cell lines (see Supplementary Figure 1a) according to standard ATCC protocols and collected cells using TRIzol reagent (Invitrogen, Waltham, MA, USA). Then, we isolated RNA using Direct-zol (Zymogen, Irvine, CA, USA). Reverse transcription was performed using the LunaScript Multiplex One-Step RT-PCR kit (New England Biolabs, Cambridge, MA) and C4A/B expression primers, as previously reported(23).

*Measure C4 protein abundance*. C4 protein abundance was measured using a capillary-electrophoresis-based system (ProteinSimple, San Jose, CA) and a commercially validated antibody directed against the alpha chain of the C4 protein (C4 Alpha Chain Polyclonal antibody; Proteintech, Rosemont, IL). Purified C4 protein was used as a positive control (CompTech, Tyler, TX, USA). Beta-actin was used as a loading control (Invitrogen, Waltham, MA, USA), and quantification of the alpha-chain peak was normalized to the actin peak.

**Results:**

*The HepG2 and Jurkat cell lines were used as positive and negative controls*, respectively. hepG2 (liver cancer cell line) produced significantly more C4A mRNA than the other cell lines tested. Among the cell lines with low C4A/B gene expression, we chose the Jurkat cell line as our negative control cell line because it is derived from immune cells (T cells). C4 protein (alpha-chain) abundance was ~67 times higher in the hepG2 cell line than in the Jurkat cell line.

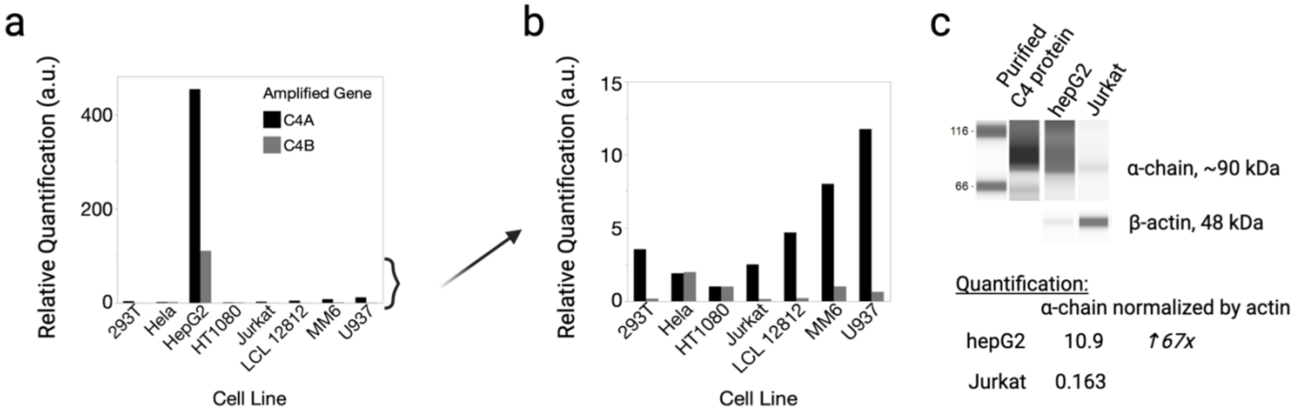


**Supplementary Figure 1: (a)** Quantitative reverse-transcription polymerase chain reaction of C4A and C4B gene expression in commonly available cell lines. The hepG2 (liver cancer cell line) produced significantly more C4A mRNA than the other cell lines tested. **(b)** Removing the hepG2 cell line, we observed the relative C4A and C4B mRNA expression in the other cell lines. Jurkat cell line was selected as the negative control cell line due to it being derived from immune cells (T cells). **(c)** ProteinSimple protein electrophoresis comparing hepG2 and Jurkat C4 protein (a-chain). Purified C4 protein was used as a positive control (CompTech, Tyler, TX, USA). Purified protein and cell line lysates were loaded based on protein quantification, however the a-chain around 90 kDa is overloaded by the close 120 kDa band (also a form of C4 protein) in the hepG2 and purified protein samples. Quantification of the a-chain (normalized by actin) indicates that the a-chain is 67 times more abundant in the hepG2 cell line compared to the Jurkat cell line.

#### B. C4A gene knockdown

To further validate that our antibody binds to the C4 protein, we created a U937 cell line with C4A/B gene knockdown.

**Methods:**

*Genetic knockdown of C4A using the CRISPR-Cas9 System*. The Gene Knockout Kit v2 was used for CRISPR-Cas9 knockdown of C4A gene (Synthego, Redwood City, CA). Single-guide RNAs (sgRNAs) selectively targeting C4A (and not C4B genes) were generated using the Synthego bioinformatics algorithm. Nucleofection was used to deliver sgRNA into the nuclei of the U937 cells (Lonza, Basel, Switzerland). After cell recovery, clonal dilution was performed to generate single-cell clonal lines according to Synthego's protocol for the clonal dilution of immortalized cell lines. Clonal lines with C4A gene knockdown were identified by protein screening. An abridged digital droplet polymerase chain reaction (determining the gene copy number of the A and B forms of the C4 gene) was performed to determine the number of copies of the C4A and C4B genes in the modified clonal lines (22).

The resulting C4A knockdown cell lines were used in flow cytometry experiments to confirm the specificity of C4 protein antibody binding. Cell lines were stained for viability (LIVE/DEAD Fixable Violet Dead Cell Stain Kit, Invitrogen, Waltham, MA) and C4 protein (Complement C4 polyclonal antibody, AbBy Fluor 647 Conjugated, Bioss, Woburn, MA). C4 protein MFI was determined only for live cells.

**Results:**

*The wild-type U937 cell line was modified to generate a U937-C4 knockdown cell line*. The single-guide RNA sequence used to target C4A for CRISPR knockdown is shown in Supplementary Figure 2a. The wild-type U937 cell line has five copies of the C4A gene (all five of which are the long form) and one copy of the C4B gene (short form). Digital-droplet PCR indicated that the U937 C4A knockdown lines had fewer copies of A and B forms of C4A (Supplementary Figure 2c). Protein screening showed a decrease in the abundance of the alpha chain (normalized to actin), with a decrease of approximately 87% after three rounds of CRISPR knockdown (in the U937$^{2A,0B}$ clonal line) compared to the *wild-type* U937$^{4A,1B}$ line (Supplementary Figure 2b). Flow cytometry indicated that the antibody targeting the C4 protein showed a 40% decrease in C4 protein intensity (as indicated by the fluorescence signal at 647 nm) in the U937$^{2A,0B}$ clonal line compared to the *wild-type* U937$^{4A,1B}$ line (Supplementary Figure 2d). We suspect a lower magnitude of downregulation due to nonspecific binding of the rabbit polyclonal antibody. Since we observed an expected decrease in the signal when we decreased the number of C4A gene copies, we conclude that both antibodies appropriately target the C4 protein.

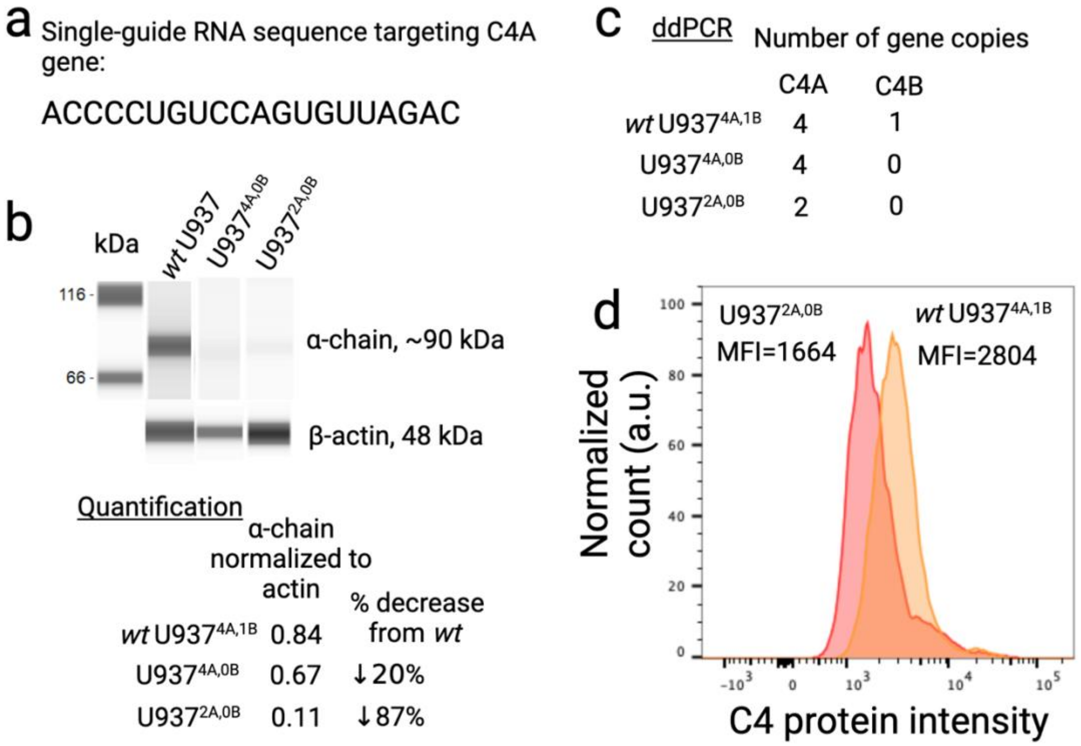


**Supplementary Figure 2:** **(a)** Single-guide RNA sequence used to target the C4A gene. **(b)** ProteinSimple protein electrophoresis comparing the *wild-type* U937 immortalized cell line to the first (U937$^{4A,0B}$) and third (U937$^{2A,0B}$) rounds of CRISPR knockdown of the C4A gene. In the third round of CRISPR knockdown, we observed a decrease in the alpha chain of the C4 protein. Quantification of the a-chain (normalized by the actin) indicates that the a-chain is decreased by approximately 87% in the third knockdown (U937$^{2A,0B}$). **(c)** Digital droplet polymerase chain reaction (ddPCR) was used to determine the number of gene copies of the C4A and C4B genes. We observed that C4B was disrupted after the first round of CRISPR. The subsequent CRISPR rounds disrupted the C4A genes, leaving only two intact C4A genes in the final clonal line (U937$^{2A,0B}$). **(d)** Fluorescence-activated cell sorting indicated that the antibody targeting the C4 protein (Complement C4 Polyclonal Antibody, AbBy Fluor 647 conjugated, bs-15186R-BF647, Bioss, Woburn, MA) showed a 40% decrease in C4 protein intensity (as indicated by fluorescence signal at 647 nm wavelength) in the U937$^{2A,0B}$ clonal line compared to the *wild-type* U937$^{4A,1B}$ line.

2. Supplementary Materials: Biobank Cohort

a

| Biobank cohort | |
|---|---|
| N of participants | 10 |
| Mean Age +/- SD | 51.6 +/- 17.9 years |
| Age range | 31 - 79 years old |
| % Male (N) | 50% (5) |
| % European Ancestry (N) | 70% (7) |
| % Black (N) | 0% (0) |
| % Asian (N) | 20% (2) |
| % Other (N) | 10% (1) |
| % Hispanic (N) | 40% (4) |

b

| Correlation between C4 protein in immune cell type and age | | |
|---|---|---|
| **Cell type** | **ρ** | **p-value** |
| Neutrophils | -0.05 | 0.89 |
| Classical monocytes | -0.07 | 0.85 |
| Nonclassical monocytes | -0.04 | 0.93 |
| Intermediate monocytes | -0.08 | 0.83 |
| NK cells | -0.03 | 0.93 |
| T cells | 0.10 | 0.78 |
| B cells | 0.10 | 0.77 |

**Supplementary Table 1: Demographics of biobank volunteers. (a)** Available demographic information on the volunteers who provided fresh whole blood samples on the day of the experiment. **(b)** Spearman's correlation between donor age and measured C4 protein levels associated with specific immune cell types.

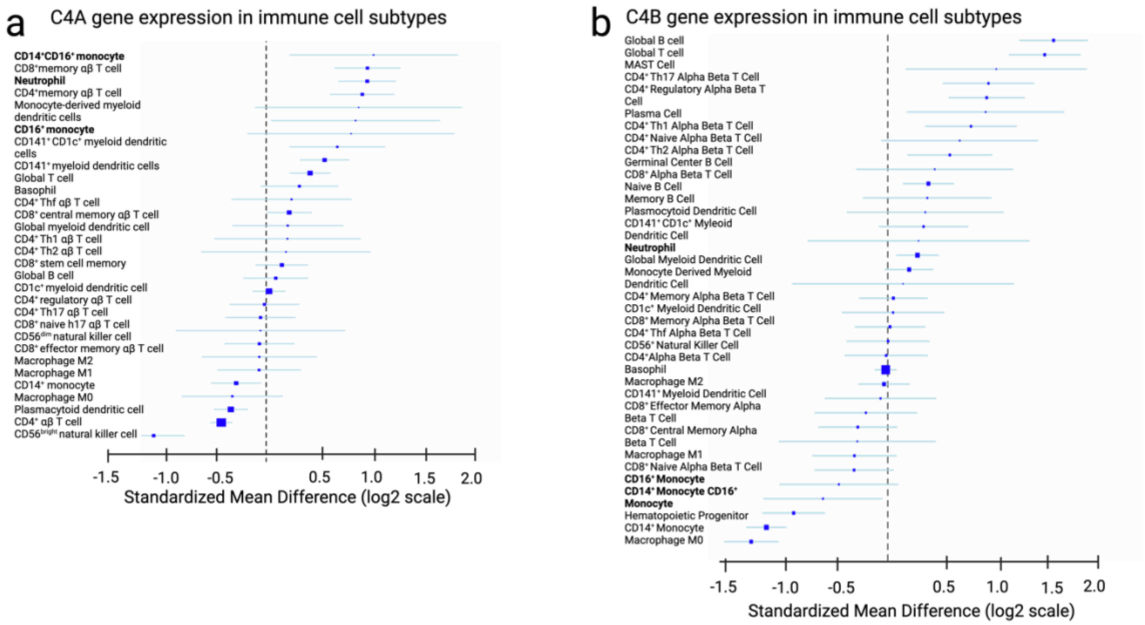


**Supplementary Figure 3: Unbiased examination of C4 gene expression in immune cell types using Meta-signature (a-b)** C4A gene expression and C4B gene expression in immune cell subtypes from Metasignature gui. More than 50,000 immune cells (and 853 available datasets) were previously analyzed in a meta-analysis of publicly available data, largely microarray data (35). Briefly, effect sizes (Hedge's g) were compared between samples from an immune cell subset of interest and all other immune cell subsets. The resulting gene expression was reported as a Standardized Mean Difference and the relative gene expression of a gene of interest in different immune cells(35). The Meta-Signature Tool (https://metasignature.stanford.edu/) was used to determine the immune cell types expressing C4A and C4B.

| Target | Clone | Conjugated Fluorophore | Manufacturer | |
|---|---|---|---|---|
| CD3 | UCHT1 | BV605 | Biolegend | San Diego, CA |
| CD14 | M5E2 | PE/Cy5 | Biolegend | San Diego, CA |
| CD16 | 3G8 | PE/Cy7 | Biolegend | San Diego, CA |
| CD19 | HIB19 | APC/Fire 810 | Biolegend | San Diego, CA |
| CD45 | HI30 | BV711 | Biolegend | San Diego, CA |
| CD56 | NCAM16.2 | BUV395 | BD Biosciences | Franklin Lakes, NJ |
| CD66b | G10F5 | FITC | Biolegend | San Diego, CA |
| HLA-DR | L243 | PE | Biolegend | San Diego, CA |

**Supplementary Table 2: Flow cytometry antibodies.** The details for the flow cytometry antibodies used for the fresh whole blood and peripheral blood mononuclear cell experiments are listed here.

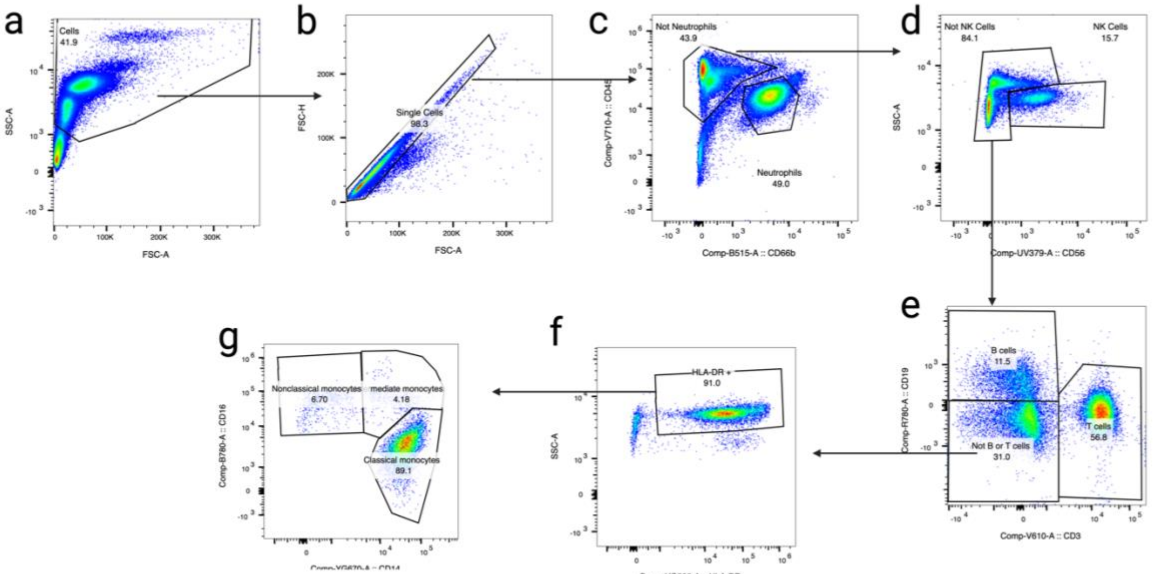


**Supplementary Figure 4: Flow cytometry gating strategy.** Flow cytometry gating of whole blood samples from donor 6 is provided as a representation of the gating strategy. Cells were gated: **(a)** first for lymphocytes based on size (area side-scatter and area forward scatter), then **(b)** for single cells by comparing the area and height of the forward scatter. **(c)** CD66b positivity and low CD45 expression in neutrophils. **(d)** Cells that were not neutrophils and were high in CD45 expression gated for Natural

Killer (NK) cells using CD56 positivity. **(e)** Cells that were not NKcells were gated for T cells ( CD3 positivity) and B cells ( CD19 positivity). **(f)** True monocytes were gated using HLA-DR expression, **(g)** CD14 and CD16 positivity for monocyte subtypes. Specifically, $CD14^{+}CD16^{-}$ cells were defined as CM, $CD14^{-}CD16^{+}$ cells as NCM, and $CD14^{+}CD16^{+}$ as Intermediate monocytes. Gates were determined using FMO controls. Compensation was performed using single fluorophore controls.

| **Comparison between immune cell types** | | **Z score** | **p-value** |
|---|---|---|---|
| Neutrophils | Classical monocytes | 3.74 | **0.0002** |
| Neutrophils | Nonclassical monocytes | 3.74 | **0.0002** |
| Neutrophils | Intermediate monocytes | 3.74 | **0.0002** |
| Neutrophils | NK cells | 3.74 | **0.0002** |
| Neutrophils | B cells | 3.74 | **0.0002** |
| Neutrophils | T cells | 3.74 | **0.0002** |
| Classical monocytes | B cells | 2.6 | **0.009** |
| Classical monocytes | T cells | 3.28 | **0.001** |
| Classical monocytes | NK cells | 1.7 | 0.09 |
| Classical monocytes | Nonclassical monocytes | 0.86 | 0.38 |
| Classical monocytes | Intermediate monocytes | 0 | 1 |
| Nonclassical monocytes | B cells | 2.38 | **0.01** |
| Nonclassical monocytes | T cells | 2.98 | **0.003** |
| Nonclassical monocytes | NK cells | 1.02 | 0.3 |
| Nonclassical monocytes | Intermediate monocytes | 1.06 | 0.29 |
| Intermediate monocytes | B cells | 2.53 | **0.01** |
| Intermediate monocytes | T cells | 3.36 | **0.0008** |
| Intermediate monocytes | NK cells | 1.87 | 0.07 |
| NK cells | T cells | 2.07 | **0.04** |
| NK cells | B cells | 1.36 | 0.17 |
| B cells | T cells | 0.64 | 0.52 |

**Supplementary Table 3: Group comparisons of C4 protein associated with different immune cells.** Flow cytometry was used to quantify the amount of C4 protein associated with distinct immune cells. The resulting C4 protein Median Fluorescent Intensity (MFI) was compared for distinct immune cell types using the Mann-Whitney U test. A comparison is presented in this figure. Comparisons between monocyte subtypes and B and T cells were omitted for clarity (clearly no difference). However, these were also included in the FDR analysis.

### 1. Supplementary Materials: Clinical Comparison Cohort

| | | Controls | Schizophrenia |
|---|---|---|---|
| Expanded Cohort | Telephone screened | 69 | 55 |
| | Consented for study | 38 | 25 |
| | Ineligible after SCID | 7 | 3 |
| | Withdrew from study | 1 | 2 |
| | Completed study | 30 | 20 |
| | Insufficient sample quantity | 1 | 0 |
| | **Total in Expanded Cohort** | **29** | **20** |
| | **Total from Pilot Cohort** | **9** | **10** |
| | **Clinical Comparison Cohort** | **38** | **30** |

**Supplementary Table 4:** The "Clinical Comparison Cohort" contains samples from the previously collected "Pilot Cohort" (22) and the "Expanded Cohort." Details of study recruitment, enrollment, and study completion of the Expanded Cohort are provided in the top portion of the table.

a

| **Neutrophil Subset** | Controls | Schizophrenia | p-value (Mann-Whitney U Test) |
|---|---|---|---|
| N of participants | 21 | 15 | |
| Mean Age +/- SD | 27.2 +/- 4.7 years | 25.7 +/- 5.9 years | 0.31 |
| % Male (N) | 48% (10) | 52% (11) | 0.39 |
| Mean BMI | 23.4 +/- 4.5 | 27.0 +/- 6.3 | 0.07 |
| % European Ancestry (N) | 48% (10) | 73% (11) | - |
| % Black (N) | 0% (0) | 7% (1) | - |
| % Asian (N) | 38% (8) | 20% (3) | - |
| % Other (N) | 14% (3) | 0% (0) | - |
| % Hispanic (N) | 14% (3) | 13% (2) | - |
| Neutrophil FST (days) | 789 +/- 300 | 632 +/- 464 | 0.46 |
| OLZ dose equivalent (mg) | N/A | 8.16 +/- 15.5 | - |

b

| **Monocyte Subset** | Controls | Schizophrenia | p-value (Mann-Whitney U Test) |
|---|---|---|---|
| N of participants | 19 | 19 | |
| Mean Age +/- SD | 25.6 +/- 5.4 years | 25.5 +/- 5.7 years | 0.86 |
| % Male (N) | 47% (9) | 53% (10) | 0.75 |
| Mean BMI | 23.4 +/- 6.7 | 26.4 +/- 6.2 | 0.08 |
| % European Ancestry (N) | 53% (10) | 58% (11) | - |
| % Black (N) | 11% (2) | 5% (1) | - |
| % Asian (N) | 37% (7) | 32% (6) | - |
| % Other (N) | 0% (0) | 11% (1) | - |
| % Hispanic (N) | 21% (4) | 11% (2) | 0.38 |
| PBMC FST (days) | 792 +/- 379 | 816 +/- 636 | 0.82 |
| OLZ dose equivalent (mg) | N/A | 9.6 +/- 14.1 | - |

**Supplementary Table 5: Demographics of study participants for neutrophil (a) and monocyte (b) subsets.** Summary statistics for Age, Sex (percentage of male participants and number of male participants), Body Mass Index (BMI), Plasma Freezer Storage Time, PBMC Freezer Storage Time and Ethnicity are provided for control and individuals with SCZ and Control groups. The distributions were compared using the Mann-Whitney U test. Abbreviations: BMI = Body Mass Index, FST = Freezer Storage Time, N = Number, OLZ = olanzapine, PBMC = Peripheral Blood Mononuclear Cells, SD = Standard Deviation.

a

| Neutrophil (Western Blot) | Controls | Schizophrenia | Whole cohort |
|---|---|---|---|
| | **Spearman's correlation of variable with measured C4 protein in CM, ρ** | | |
| Age | 0.46, p = 0.04 | 0.09, p = 0.76 | 0.26, p = 0.12 |
| Body Mass Index | 0.40, p = 0.07 | 0.31, p = 0.26 | 0.26, p = 0.12 |
| PBMC FST (days) | -0.20, p = 0.37 | 0.35, p = 0.20 | 0.11, p = 0.54 |
| OLZ dose equivalent (mg) | NA | 0.23, p = 0.42 | -0.16, p = 0.36 |
| Number of C4A gene copies | 0.05, p = 0.84 | 0.63, **p = 0.01** | 0.25, p = 0.14 |
| Number of C4B gene copies | 0.09, p = 0.70 | -0.12, p = 0.67 | -0.05, p = 0.77 |
| **Compare means of reported participant sex with measured C4 protein in CM, p-value** | | | |
| Reported Biological Sex | p = 0.97 | p = 0.76 | p = 0.72 |
| Female | 0.16 +/- 0.06 | 0.15 +/- 0.11 | 0.16 +/- 0.08 |
| Male | 0.20 +/- 0.13 | 0.14 +/- 0.08 | 0.17 +/- 0.11 |

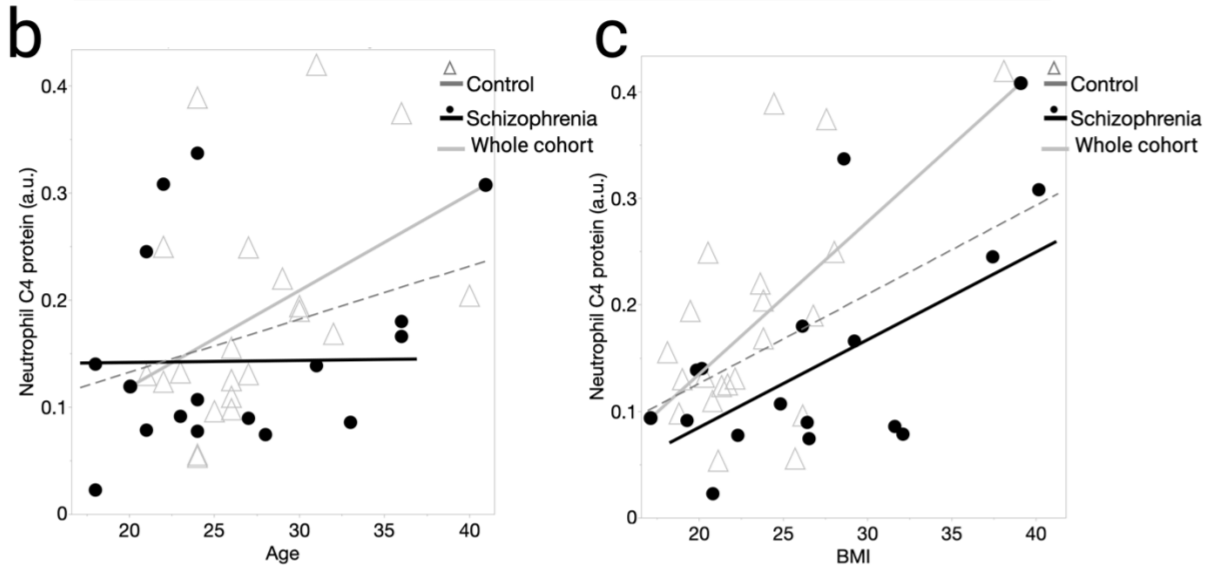


**Supplementary Figure 5: Examining the role of potential confounders in measured neutrophil C4 protein abundance. (a)** Table showing the analysis results Spearman correlation of neutrophil C4 protein abundance from flow cytometry and age, body mass index, PBMC freezer storage time (FST) in days and antipsychotic medication in olanzapine dose equivalents (OLZ dose equivalent) for the control group, SCZ group and the combined (control and SCZ) 'whole' cohort (except for OLZ dose equivalents which was only performed for the patient group). **(b-c)** Linear regression between neutrophil C4 protein abundance and **(b)** age and **(c)** body mass index (BMI).

a

| Classical monocyte (Flow Cytometry) | Controls | Schizophrenia | Whole cohort |
|---|---|---|---|
| | **Spearman's correlation of variable with measured C4 protein in CM, ρ** | | |
| Age | 0.09, p = 0.58 | 0.22, p = 0.25 | 0.18, p = 0.14 |
| Body Mass Index | -0.16, p = 0.33 | -0.21, p = 0.27 | -0.22, p = 0.07 |
| PBMC FST (days) | -0.26, p = 0.12 | -0.30, p = 0.10 | -0.32, **p = 0.01** |
| OLZ dose equivalent (mg) | NA | 0.13, p = 0.52 | -0.13, p = 0.26 |
| Number of C4A gene copies | -0.18, p = 0.30 | -0.13, p = 0.49 | -0.17, p = 0.18 |
| Number of C4B gene copies | -0.03, p = 0.88 | 0.09, p = 0.62 | 0.01, p = 0.92 |
| **Compare means of reported participant sex with measured C4 protein in CM, p-value** | | | |
| Reported Biological Sex | p = 0.89 | p = 0.71 | p = 0.89 |
| Female | 2.54 +/- 2.15 x $10^4$ | 1.69 +/- 1.37 x $10^4$ | 2.22 +/- 1.90 x $10^4$ |
| Male | 2.62 +/- 1.87 x $10^4$ | 1.72 +/- 1.87 x $10^4$ | 2.19 +/- 1.91 x $10^4$ |

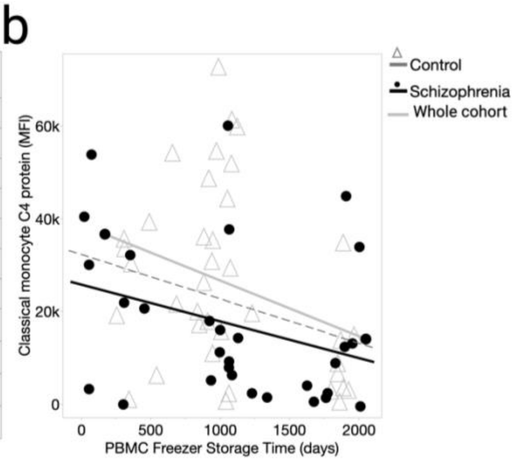


c

| Classical monocyte (Immunofluorescence) | Controls | Schizophrenia | Whole cohort |
|---|---|---|---|
| | **Spearman's correlation of variable with measured C4 protein in CM, ρ** | | |
| Age | 0.18, p = 0.47 | 0.14, p = 0.56 | 0.14, p = 0.39 |
| Body Mass Index | 0.17, p = 0.49 | 0.23, p = 0.34 | 0.15, p = 0.38 |
| PBMC FST (days) | -0.02, p = 0.93 | 0.46, **p = 0.05** | 0.32, **p = 0.05** |
| OLZ dose equivalent (mg) | NA | 0.36, p = 0.13 | -0.13, p = 0.43 |
| Number of C4A gene copies | -0.20, p = 0.42 | -0.14, p = 0.57 | -0.20, p = 0.22 |
| Number of C4B gene copies | -0.02, p = 0.94 | 0.23, p = 0.35 | 0.07, p = 0.66 |
| **Compare means of reported participant sex with measured C4 protein in CM, p-value** | | | |
| Reported Biological Sex | p = 0.78 | p = 0.97 | p = 1.00 |
| Female | 1.65 +/- 0.66 x $10^4$ | 1.34 +/- 0.65 x $10^4$ | 1.51 +/- 0.65 x $10^4$ |
| Male | 1.66 +/- 0.74 x $10^4$ | 1.50 +/- 0.97 x $10^4$ | 1.57 +/- 0.85 x $10^4$ |

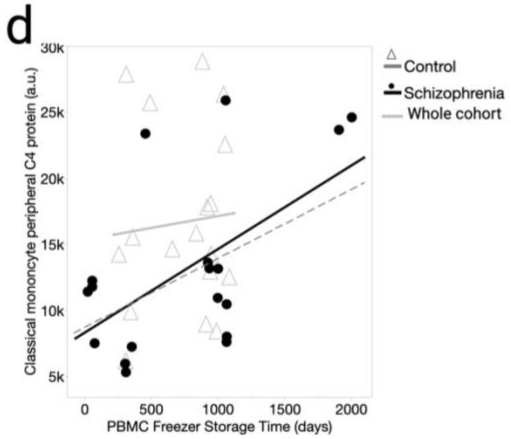


**Supplementary Figure 6: Examining the role of potential confounders in measured classical monocyte C4 protein abundance.** **(a)** Table showing the analysis results Spearman correlation of CM C4 protein abundance from flow cytometry and age, body mass index, PBMC freezer storage time (FST) in days and antipsychotic medication in olanzapine dose equivalents (OLZ dose equivalent) for the control group, SCZ group and the combined (control and SCZ) 'whole' cohort (except for OLZ dose equivalents which was only performed for the patient group). **(b)** Linear regression between flow cytometry-determined C4 protein abundance in CM and PBMC freezer storage time. **(c)** Table showing the analysis results Spearman correlation of CM C4 protein abundance from immunofluorescence and age, body mass index, PBMC freezer storage time (FST) in days and antipsychotic medication in olanzapine dose equivalents (OLZ dose equivalent) for the control group, SCZ group and the combined (control and SCZ) 'whole' cohort (except for OLZ dose equivalents which was only performed for the patient group). **(d)** Linear regression between Immunofluorescence-determined C4 protein abundance in CM and PBMC freezer storage time for the control group, SCZ group and the combined (control and SCZ) 'whole' cohort.
